# Paternal multigenerational exposure to an obesogenic diet drives epigenetic predisposition to metabolic disorders

**DOI:** 10.1101/2020.05.19.104075

**Authors:** Georges Raad, Fabrizio Serra, Luc Martin, Marie-Alix Derieppe, Jérôme Gilleron, Vera L Costa, Didier F Pisani, Ez-Zoubir Amri, Michele Trabucchi, Valérie Grandjean

## Abstract

Obesity is a growing societal scourge responsible for approximately 4 million deaths worldwide. Recent studies have uncovered that paternal excessive weight induced by an unbalanced diet affects the metabolic health of offspring. These reports mainly employed single-generation male exposure. However, the consequences of multigenerational unbalanced diet feeding on the metabolic health of progeny remain largely unknown. Here, we show that maintaining paternal western diet feeding for five consecutive generations in mice induces a gradual enhancement in fat mass and related metabolic diseases over generations. Strikingly, chow-diet-fed progenies from these multigenerational western-diet-fed males develop a “healthy” overweight phenotype that is not reversed after 4 subsequent generations. Mechanistically, sperm RNA microinjection experiments into zygotes suggest that sperm RNAs are sufficient for establishment but not for long-term maintenance of epigenetic inheritance of metabolic pathologies. Progressive and permanent metabolic deregulation induced by successive paternal western-diet-fed generations may contribute to the worldwide epidemic of metabolic diseases.

## Introduction

Nongenetic inheritance of newly acquired phenotypes is a new concept in biology whereby changes induced by specific environmental cues in parents (mothers and/or fathers) can be transmitted to the next generation [1-3]. This process is evolutionarily conserved and has been described from worms to humans [4-7]. The fact that environmental cues have the potential to modify the molecular hereditary information carried by the spermatozoa demonstrates that the environmentally induced epigenetic modifications [8][8] are not erased through the epigenetic reprogramming process, causing them to be inherited by the next generations [9, 10]. Although the role of epigenetic modifications such as DNA methylation [9, 11, 12] and chromatin modification [13, 14] cannot be excluded in this process, independent experimental data strongly evoke the central role of sperm RNA as a vector of paternal intergenerational epigenetic inheritance of, at least, environmentally induced metabolic pathologies [1, 3, 15]. Unlike genetic inheritance, environmentally induced epigenetic alterations are reversible, enabling the loss of previously acquired characteristics [16]. Although environmental changes might persist over several generations, most reports have been based on the maintenance of paternal environmental cues for just one generation [17]. This is particularly true for certain lifestyle habits, such as eating high-fat and high-sugar junk food, also called a Western diet (WD). Thus, although people around the world may face multigenerational unbalanced nutrition, there have been limited studies on its effects on the metabolic health of the progeny.

Herein, we studied the impact of the paternal maintenance of an unhealthy WD for multiple generations on the metabolic phenotype of both the progenitors and their respective chow-diet-fed (CD-fed) offspring.

## Results

### Feeding successive paternal generations with a diet exacerbates the overweight phenotype and accelerates the development of obesity-associated pathologies

To test experimentally whether the maintenance of an unhealthy diet through the paternal germline influences the metabolic phenotype of the resulting individuals, C57BL6/J male mice were fed a WD for five consecutive generations (from WD1 to WD5) (**Fig 1A**). According to a previous study [18], the average body weight of the WD-fed male mice increased gradually with multigenerational WD feeding **(Fig 1B** and **S1A Fig)**. This gradual increase in total body weight with paternal multigenerational WD feeding was associated with a gradual increase in perigonadal white adipose tissue (gWAT) mass (**Fig 1C**). Indeed, the gWAT volume measured by computed tomography increased 2.3-fold and 3.4-fold in WD1 and WD5 mice, respectively, compared to that of control mice (CD-fed mice) (**S1 Table**). The increase in gWAT mass was positively correlated with total body weight (perigonadal fat mass versus total body weight; Spearman’s *r* =0.78, *p* < 0.0001, **S1B Fig**). It was also associated with the hypertrophy of white adipocytes, with a median surface cell area of white adipocytes increasing from 1500 to 4000 μm^2^ from the first (WD1) to the fifth generation (WD5) and with a decreased calculated number of adipocytes in WD5 compared to the controls (**Fig 1D-1F**). Furthermore, our RNA-seq comparison between the gWAT of WD1 and WD5 males revealed that multigenerational WD feeding has a strong impact on the gWAT gene expression profile. In fact, we observed an increase in differentially expressed genes (DEGs), from 325 in WD1 (with 93 upregulated and 232 downregulated genes) to 1199 (757 upregulated and 442 downregulated) in WD5, compared to the respective CD-fed mice. Interestingly, while the majority of DEGs in WD1 (66%) were also deregulated in WD5, a minority of DEGs in WD5 (only 8% for the upregulated genes and 35% for the downregulated genes) were deregulated in WD1 (p value<0.01). Importantly, all common genes were deregulated in the same direction (**Fig 1G-1H**). Interestingly, querying the WD1 and WD5 DEGs against the molecular signature database (MSigDB) collection of curated gene pathway annotations revealed a specific WD5 enrichment in gene sets associated with CHEN_METABOLIC_SYNDROM_NETWORK (genes forming the macrophage-enriched metabolic network (MEMN) claimed to have a causal relationship with metabolic syndrome traits) and with genes potentially regulated by the methylation of lysine 4 (H3K4) and lysine 27 (H3K27) of histone H3 and by polycomb repressive complex 2 (PRC2) **(S2A-S2B Table 2)** [19].

**Fig 1.**
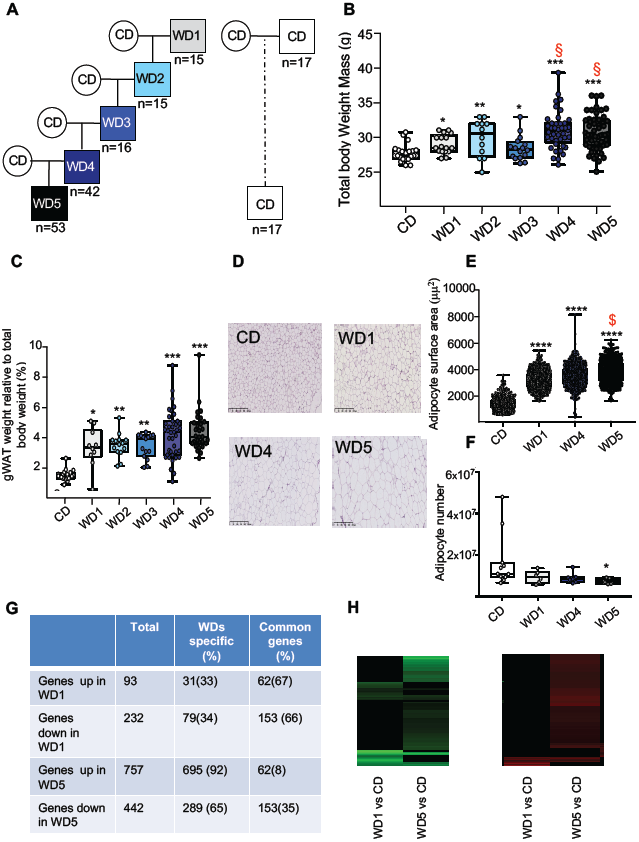
Five consecutive paternal generations of WD feeding exacerbate the WD-induced overweight phenotype. (A) Study design for the maintenance of WD feeding for 5 consecutive generations through the paternal lineage. Male mice were randomized to receive either a control diet (CD; 5% of energy from fat) or a high-fat diet (WD1; 45% of energy from fat) for 3 months before being mated with CD-fed females to generate WD2 offspring. Five independent WD2 males fed a WD for 3 months were mated with CD-fed females to generate WD3 offspring. The same procedure was repeated twice to generate the WD4 and WD5 offspring. (B) Box-whiskers (min- max) of the median total body weight of the different male WD cohorts (n≥8 mice per group). (C) Box-whiskers (min-max) of the median perigonadal white adipose tissue (gWAT) weight relative to total body weight in the different WD cohorts. (D) H&E staining of gWAT sections (scale bar: 200 μm) in representative CD, WD1, WD4 and WD5 males. (E) Box-whiskers (min- max) of the median surface area (μm^2^) of the adipocytes, which was calculated using Image Analyzer software (ImageJ). The total count ranged from 3275 to 7052 cells per condition (n≥4 mice per group). (F) Box-whiskers (min-max) of the number of adipocytes which was estimated using the mathematical equation developed by Jo et al.[32], as previously described in [20]. **g**, Table showing the differentially expressed genes (DEGs) in WD1 and WD5 perigonadal white adipose tissue. (H) Heatmap diagrams of DEGs (p<0.01 log2FC≥|0.6|) in both WD1 and WD5 perigonadal white adipose tissue compared to expression in the CD gWAT tissue cohort. Negative log-ratios (log fold change) are shown in green, while positive log-ratios are shown in red. Genes that are differentially expressed in both WD1 and WD5 are deregulated in the same way (n=3 mice/group). *p<0.05, **p<0.01, ***p<0.001 (the Kruskal-Wallis test, a rank-based nonparametric test for multiple comparisons, two-stage linear step-up procedure of Benjamin, Krieger and Yekutieli was used to calculate the adjusted *p* value). § denotes the WD groups whose median was significantly different from that of the WD1 cohort.

The aforementioned modulations of white adipose tissue in WD generations shed light on the possible exacerbation of obesity-associated pathologies (such as insulin resistance (and subsequently type-II diabetes) and nonalcoholic fatty liver disease) [20]. To check this hypothesis, several metabolic risk parameters related to these pathologies were analyzed in WD-fed mice (**Table 1**). In comparison with CD-fed mice, circulating plasma levels of leptin, C-reactive protein (CRP), one marker of inflammation, and total cholesterol were significantly higher in the WD3 (p<0.01), WD4 (p<0.05) and WD5 (p<0.01) groups but not in the WD1 (p=0.07) or WD2 (p=0.4) groups (**Table 1**). The gradual alterations in these metabolic parameters over generations were found to be positively correlated with the increase in gWAT mass (**S1C-S1E Fig**). At the molecular level, the progressive increase in serum leptin over WD-fed generations was positively correlated with a gradual increase in leptin mRNA levels in the gWAT of the respective male mice (total plasma leptin and *leptin* mRNA, Spearman’s *r* =0.89, *p* < 0.0001, **S1F Fig**), suggesting an accumulation of epigenetic modifications of the leptin promoter. These results are in line with recent studies showing that leptin upregulation occurs via epigenetic malprogramming in white adipose tissue [21, 22]. Furthermore, we found a significantly impaired response in the intraperitoneal glucose tolerance test (GTT) in all WD-fed mouse groups (**Fig 2A**), which was not associated, except for in WD2-fed males, with an impaired insulin response, as shown by the intraperitoneal insulin tolerance test (ITT) (**Fig 2B**). Therefore, unlike the other metabolic parameters, we did not notice any significant exacerbation of insulin sensitivity in successive generations. Moreover, the response to an intraperitoneal glucose tolerance test (measured through the AUC-GTT calculation) was not correlated with the gWAT mass **(S1G Fig**). Together, these data might reflect the multifactorial and complex nature of the pathogenesis of obesity-induced diabetes.

**Table 1.** Evolution of serum biomarker parameters in different WD groups.

| Parameters | Control<br>n=6 | WD1<br>n=4 | WD2<br>n=5 | WD3<br>n=7 | WD4<br>n=7 | WD5<br>n=7 |
| --- | --- | --- | --- | --- | --- | --- |
| Adiponectin (μg/ml) | 44.31±10 | 51.16±20 | 37.30±9 | 40.25±15 | 3542±698 | 43.90±11 |
| Leptin (μg/ml) | 6.4±1.6 | 11.1±4.5 | 9.2±3.9 | 15.8.55±9.75 | **19.9±9.98\*\*\$** | **17±19.8\*\*\$** |
| CRP (ng/ml) | 3896±1223 | 5694±585 | 5787±459 | **7723±2050\*\*** | **5813±1840\*** | **6567±1036\*\*** |
| Total cholesterol (mg/dl) | 0.9±0.2 | **1.5±0.4\*\*** | 1.33±0.39 | **1.8±0.2\*** | **1.8±0.3\*\*\*\$** | **1.4±0.33\*\*\*\$** |

Values are expressed as the mean ± SD. Kruskal-Wallis test, a rank-based nonparametric test for multiple comparisons, two-stage linear step-up procedure of Benjamin, Krieger and Yekutieli was used to calculate the *p* value. Numbers are in bold if *p*<0.05. \$ denoted the WD groups whose median was significantly different from that of the WD2 group. \**p*<0.05, \*\**p*<0.01.

**Fig 2.**
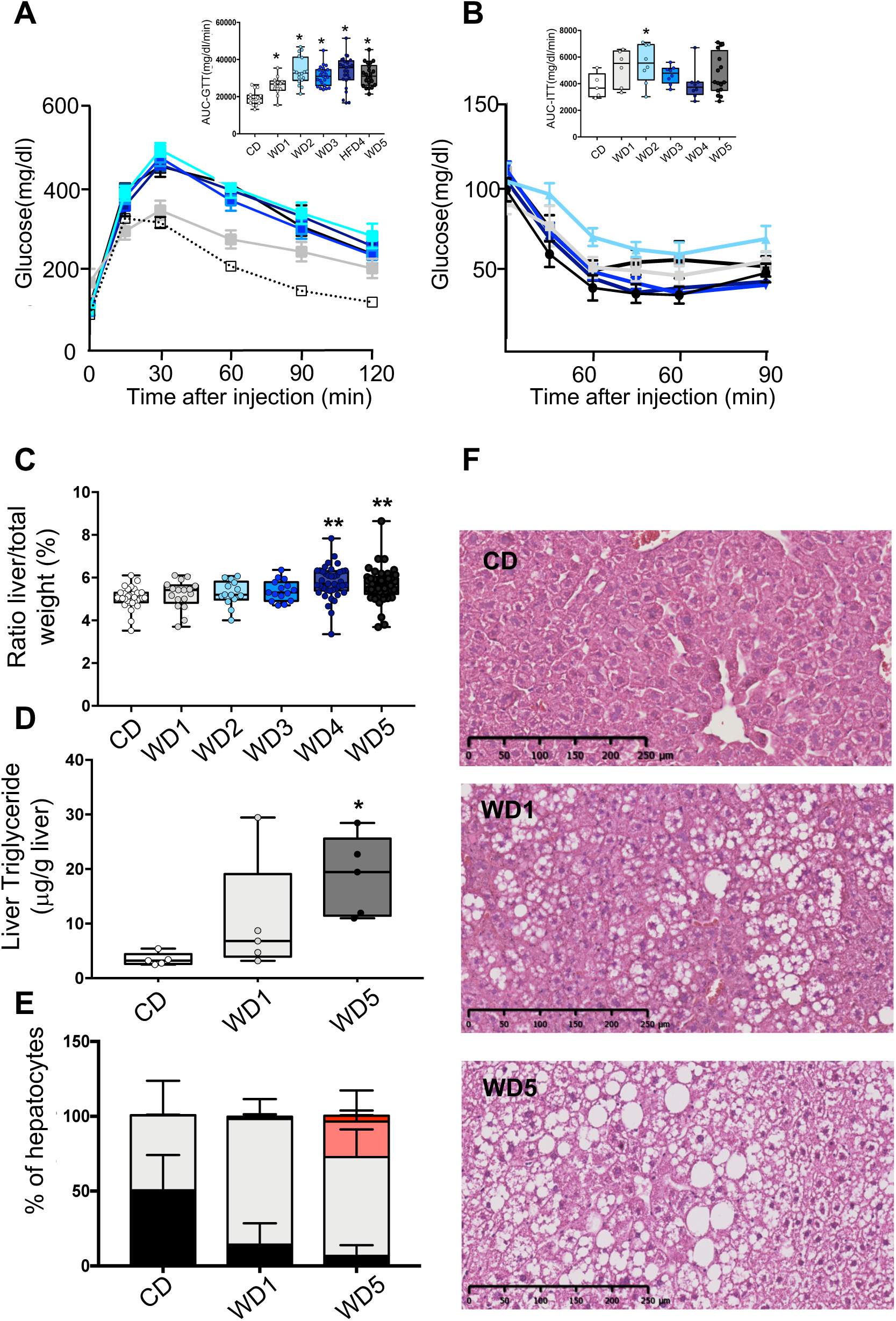
Five consecutive paternal generations of WD feeding exacerbate WD-induced overweight pathologies. (A-B) Evolution of glucose parameters in male mice fed a WD for five successive generations. Blood glucose and insulin tolerance tests were performed on 16-week-old males (n≥6). Plasma glucose [inserted box-whiskers (min-max) of the median area under the curve (AUC) and above baseline for glucose from time point 0 to 120; glucose tolerance test] (A) [inserted box- whiskers (min-max) of the median AUC and above baseline for glucose from time point 0 to 100; insulin tolerance test] (B). Glucose tolerance and insulin tolerance tests were conducted in the morning in overnight-fasted mice. **c** Box-whiskers (min-max) of the median liver weight relative to total body weight in the different WD cohorts (n≥8 mice per group). (D) Liver triglyceride contents in the CD, WD1 and WD5 groups (n≥6). (E) Percentage of normal hepatocytes (black boxes), hepatocytes with microvesicular steatosis (gray boxes) and hepatocytes with macrovesicular steatosis (pink boxes) in CD, WD1 and WD5 livers (n≥6). (F) H&E staining of liver sections (scale bar: 250 μm) from representative CD, WD1 and WD5 males. *p<0.05, **p<0.01, ***p<0.001 (the Kruskal-Wallis test, a rank-based nonparametric test for multiple comparisons, two-stage linear step-up procedure of Benjamin, Krieger and Yekutieli was used to calculate the adjusted *p* value). § denotes the WD groups whose median was significantly different from that of the WD1 cohort.

Strikingly, although the C57BL6/J-strain male mice fed a WD diet for one generation failed to develop strong alterations in liver phenotype [23, 24], major abnormalities were observed in WD5 liver, i.e., organ weight, histological and biochemical parameters. Indeed, the mass of the WD5 liver (not that of the WD1 liver) was significantly higher than that of the CD specimens **(Fig 2C)**. Furthermore, unlike WD1 liver, histological and biochemical examinations revealed the presence of macrovesicular steatosis with significantly increased triglyceride (TG) levels in WD5 liver compared with CD liver (p<0.01, respectively) **(Fig 2D-F)**. Therefore, the phenotype of WD5 livers exhibits typical features of fatty liver.

Together, both morphological and molecular features demonstrate that multigenerational WD feeding induced a progressive dysregulation of the male metabolic phenotypes (**S1H Fig**), with an exacerbation of the gWAT size and gWTA transcriptional alteration as well as of obesity-associated pathologies such as fatty liver. Therefore, a worsening of the underlying medical conditions can be potentially transmitted to next generations.

### Long-term transgenerational epigenetic inheritance of an overweight “healthy” phenotype

Previous reports showed that WD-induced metabolic dysregulations during one-generation exposure could be transmitted across 1 (F1) or 2 generations (F2) fed a CD^2 10^. To investigate the impact of feeding a WD through several generations on the inheritance of diet-induced metabolic pathologies, we compared the metabolic status of F1, F2 and F3 cohorts fed a CD generated from either WD1 or WD5 males (**Fig 3A**). As expected from previous studies [1-3], male and female F1 progenies derived from WD1 males (F1-WD1) were heavier than the control animals with CD-fed ancestors (**Fig 3B and 3F**). Although the difference did not reach significance at the age of 18 weeks, the same trend was also observed for the F1 progenies derived from WD5 (F1-WD5) male progenies (**Fig 3D and 3H**). This overweight phenotype was associated with impaired glucose tolerance as measured by the GTT for both the male F1-WD1 and F1-WD5 progenies and the female F1-WD5 mice **(S2E and S2G Fig and S3-S4 Tables**). We noticed, however, the absence of intergenerational inheritance of the fatty liver phenotype observed in the WD5 progenitors.

**Fig 3.**
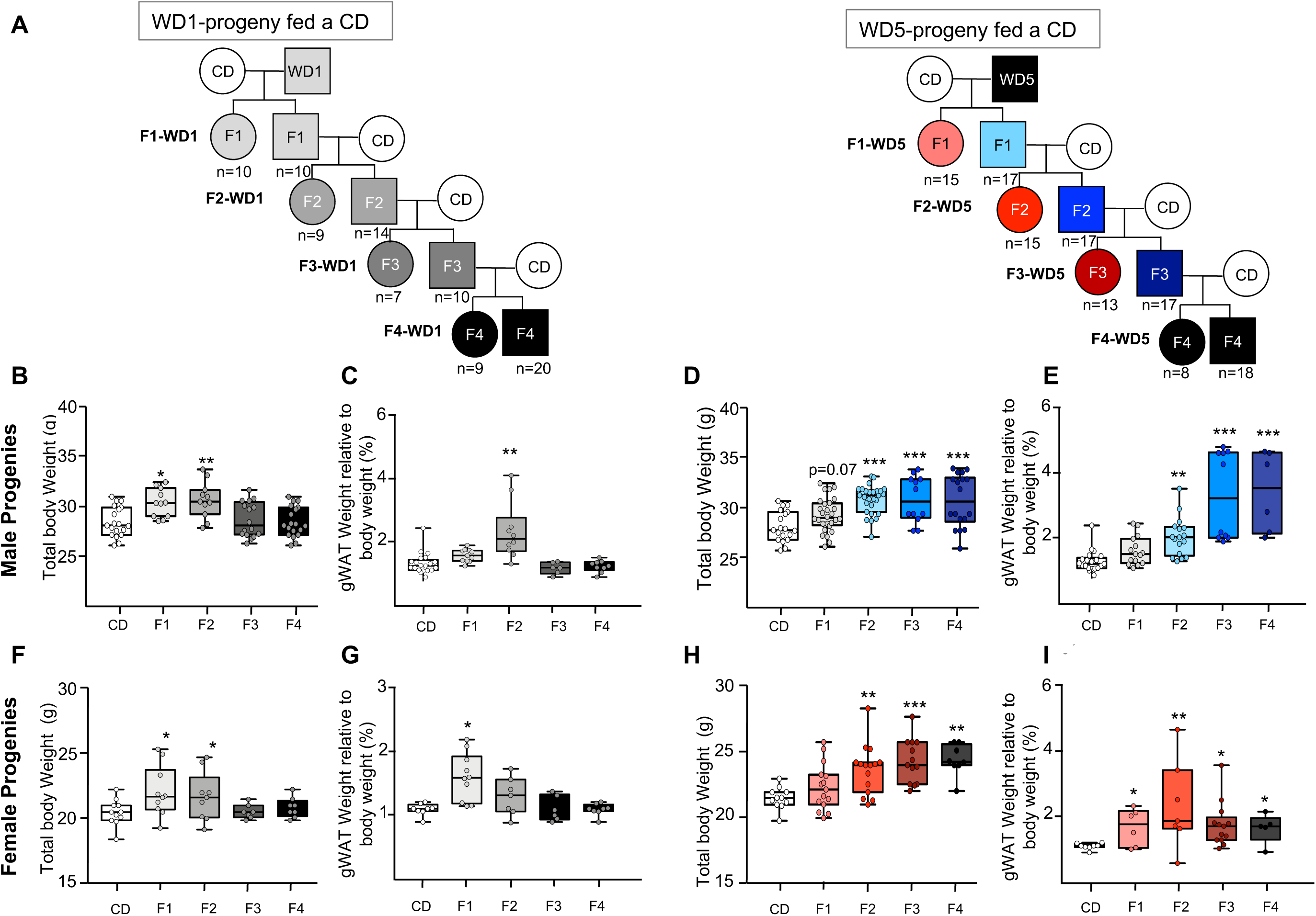
Maintenance of the overweight phenotype after 4 generations on the CD in the progenies generated from WD5-fed males. (A) Study design for the inheritance of WD-induced metabolic alterations in WD1- and WD5- fed animals. Five WD1 and five WD5 male mice from different littermates fed the control diet (CD) were mated with CD-fed females to generate F2-WD1 and F2-WD5 offspring, respectively. Each offspring was fed the CD. This crossing scheme was repeated twice to obtain the F3-, F4-WD1 and F3-, F4-WD5 offspring. The number of mice is indicated. Box-whiskers (min-max) of the median total body weights of 18-week-old males (B, D) and females (F, H) of progenies from WD-fed animals. Box-whiskers (min-max) of the median gWAT of males (C, E) and females (G, I) of progenies from WD-fed animals. Gray rectangles represent the male and female progenies from WD1-fed animals. Blue and red dots represent the male and female cohorts, respectively, of progenies from WD5-fed animals. *p<0.05, **p<0.01, ***p<0.001 (the Kruskal-Wallis test, a rank-based nonparametric test for multiple comparisons, two-stage linear step-up procedure of Benjamin, Krieger and Yekutieli was used to calculate the adjusted *p* value).

Both male F2-WD1 and F2-WD5 CD-fed progenies were also overweight (p<0.01). This phenotype was associated with an excessive accruement of gWAT mass of at least 90% over the control (**Fig. 3C-3E-3G-3I**). Importantly, although the female and male F2-WD5 progenies were found to be significantly fatter and heavier than the F2-WD1 cohorts, these mice did not exhibit impaired glucose tolerance (as measured by the GTT) **(S2E, S2G Fig)** or signs of fatty liver lesions **(S3 Fig).**

The metabolic differences were even more striking in both F3 and F4 progenies **(S2A-2D Fig)**. Thus, while the F3-WD1 progenies exhibited metabolic characteristics very similar to control mice, both males and females of the F3-WD5 progenies were significantly heavier and fatter (p<0.001 and p<0.01, respectively) than control mice (**Fig 3B-3I** and **S3 and S4 Tables)**. In parallel, the overweight phenotype was associated only with an increase in gonadal fat mass, which persisted in the F4-WD5 progenies **(Fig 3)**. Strikingly, despite being overweight, the progenies derived from WD5-fed animals did not display any alteration in terms of glucose metabolism **(S2E, S2F, S2G, S2H Fig)** and fatty liver pathologies at 4 months of age **(S3 Fig) (S3 and S4 Tables**).

Collectively, these data suggest that WD feeding for multiple generations induces stable germline epigenetic modifications that were not erased after removing the stressor(s) for at least 4 generations of CD-fed progeny.

### Sperm RNAs transmit only transient epigenetic inheritance of WD-induced pathologies

Specific signatures of sperm small RNAs from WD-fed mice have been previously shown to act as a vector of intergenerational epigenetic inheritance of newly acquired pathologies [1, 3, 15, 25]. To determine whether sperm small RNAs are also involved in the long-term maintenance of epigenetic inheritance (transgenerational epigenetic inheritance), we first searched for small RNA DEGs (adjusted p value<0.05) between WD (WD1 or WD5) and CD sperm. As shown in **S5 and S6 Tables**, we identified 584 and 614 DEGs in WD1 and WD5, respectively, compared to the control mice. Interestingly, approximately one-third of DEGs (190 sequences) were present in both WD1 and WD5 RNA sperm populations **(S4 Fig)**. Among these common small RNAs, we identified several tRNA fragments and microRNAs known to be involved in short-term epigenetic inheritance of metabolic dysfunction (intergenerational inheritance) [1, 3] **(S4D-S4E Fig).** These data indicate that sperm RNAs could be involved in the epigenetic inheritance of metabolic alterations in both WD1 and WD5 males.

To further investigate the role of sperm RNAs in the long-term transgenerational epigenetic inheritance of metabolic alterations, microinjection experiments into naïve zygotes were performed with total sperm RNA from either WD1 or WD5 males (RNA-WD1 progenies and RNA-WD5 progenies, respectively) (**Fig 4A)**. As previously reported, this experiment faithfully reproduces the pattern of short-term paternal transmission of environmentally induced phenotypes in crosses[1, 3, 4, 15, 25]. In agreement with previous studies, male 12-week F1- RNA-WD1 and F1-RNA-WD5 progenies were heavier than F1-RNA-CD progenies (31 g vs 30 g, p<0.05) (**Fig 4B)**. In addition, they displayed glucose and insulin response alterations, as shown by GTT and ITT analyses, with significantly higher values of the area under the curve than the controls (**Fig 4D, 4E** and **S7 Table).** Regarding the fatty liver phenotype, neither abnormal TG levels nor histological abnormalities were observed in livers from F1-RNA-CD and F1-RNA-WD progenies. Thus, the metabolic alterations observed in F1-RNA progenies are partially reminiscent of the WD1 and WD5 male phenotype.

**Fig 4.**
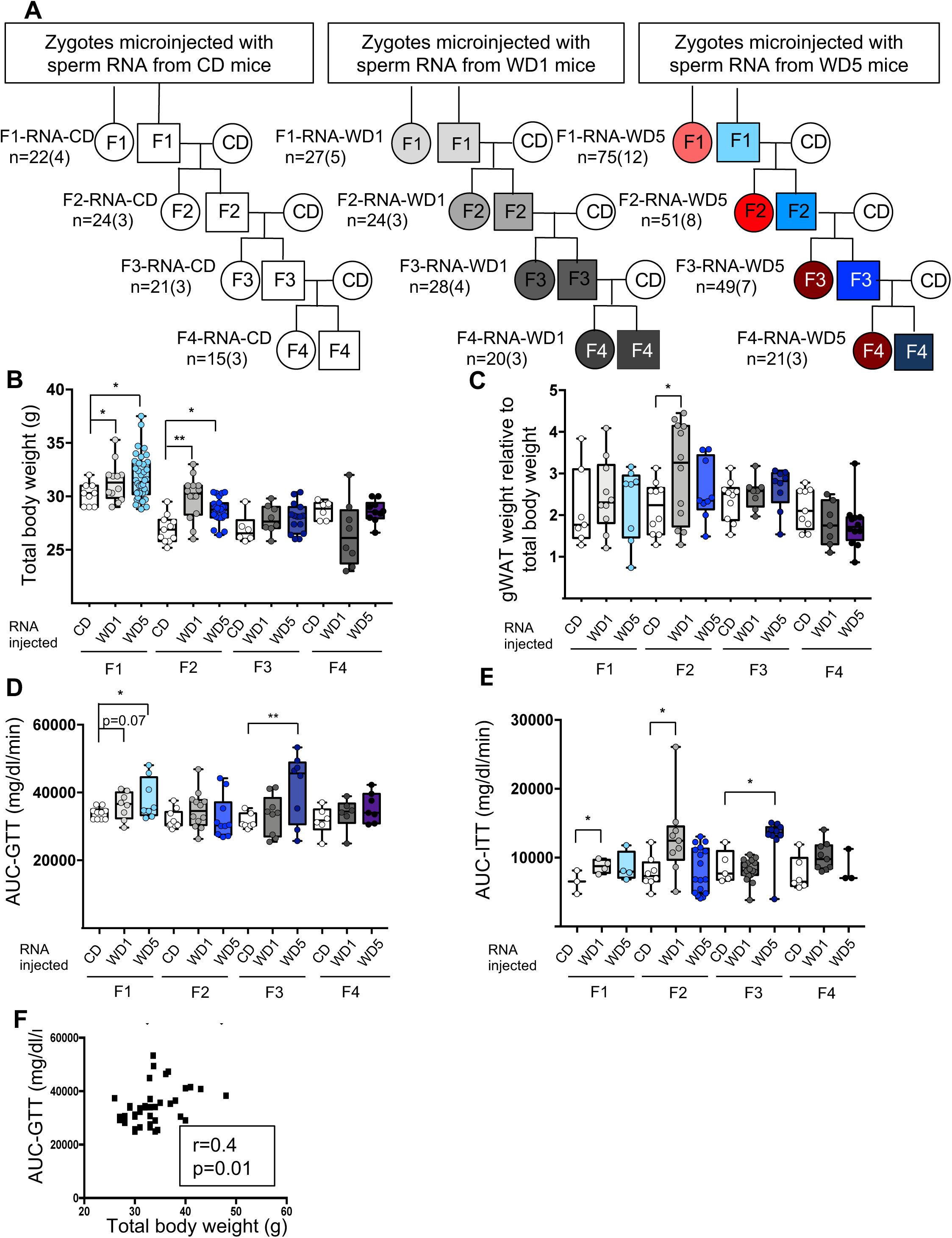
Zygotic microinjection of sperm total RNA from either WD1 or WD5 males induces metabolic alterations in the F1 and F2 CD-fed progenies that are not maintained in the F3 and F4 CD-fed progenies. (A) Study design for the inheritance of metabolic alterations induced after the microinjection of sperm total RNA from CD-, WD1- or WD5-fed males into C57BL/6J zygotes. Five F1 CD-fed males from each set of RNA microinjections were mated with CD-fed females to generate F2- RNA offspring. Each offspring was fed a control diet. This crossing scheme was repeated twice to obtain the F3-RNA offspring and then the F4-RNA offspring. (B) Box-whiskers (min-max) of the median total body weight of the F1-, F2-, F3-, and F4-RNA male progenies (n≥8 mice per group). (C) Box-whiskers (min-max) of the median gWAT weight relative to total body weight in the different RNA progenies. The evolution of glucose parameters in male mice from RNA-injected progenies. (D) Box-whiskers (min-max) of the median AUC-GTT of each cohort. (E) Box-whiskers (min-max) of the median AUC-ITT of each group. (F) Bivariate correlation between the body weight of the F2-RNA-CD, F2-RNA-WD1 and F2-RNA-WD5 progenies and the AUC-GTT (n=38). This correlation was similar using parametric (Pearson, *r* = 0.4, *p* = 0.01) or nonparametric (Spearman, *r* = 0.4, *p* = 0.01) correlations. *p<0.05, **p<0.01, ***p<0.001 (the Kruskal-Wallis test, a rank-based nonparametric test for multiple comparisons, two-stage linear step-up procedure of Benjamin, Krieger and Yekutieli was used to calculate the adjusted *p* value). § denotes the WD groups whose median was significantly different from that of the WD1 group.

Overweight phenotypes and glucose response alterations were partially transmitted to the F2 and F3 generations (**Fig 4, S8 and S9 Tables)**. Intriguingly, while we did not observe any liver abnormalities in F1-RNA progenies, liver histological examinations revealed macro- and microvesicular steatosis in hepatocytes of two F2-WD overweight males (2 out of 10) **(S5 Fig)**. It should be noted that these abnormal hepatocytes were never observed in RNA-CD progenies. Nevertheless, all the metabolic alterations were completely absent in the F4 generations (**Fig 4 and S10 Table**).

The metabolic observed phenotype of WD1 and WD5 progenies obtained by either RNA microinjection or natural mating exhibited some discrepancies. First, the overweight phenotype induced by sperm RNA from WD5 males was not exacerbated compared to that induced by sperm RNA from WD1 males. In fact, no statistically significant difference was observed among the body weights of the F1, F2 and F3 progenies derived from sperm RNA of WD1- and WD5 animals. Second, the sperm-RNA-induced overweight phenotype was associated with glucose metabolic alterations (total body weight and AUC-GTT, Spearman’s *r* =0.4, *p* < 0.01, **Fig 4F**) and was sporadically associated with fatty liver abnormalities, in both WD1 and WD5 **(S5 Fig)**. Last, in contrast to natural mating of WD5, the sperm-RNA-induced overweight phenotype was not transgenerationally inherited, while the metabolic abnormality was very stable in the progenies obtained from natural mating of WD5. Taken together, these data strongly suggest that sperm RNAs are not sufficient for the long-term epigenetic inheritance of metabolic dysfunctions.

## Discussion

Growing evidence suggests that an unbalanced diet of the father negatively affects its metabolic health and that of its progenies. Of particular interest, little attention has been focused on the effect of paternal successive generations of unbalanced diet exposure on metabolic health, which may have public health and economic impacts. To this end, we fed male mice for 5 successive generations on a high-fat, high-sugar diet (western diet, WD) to compare the metabolic parameters across multiple generations of WD males and to assess the persistence of the WD-induced metabolic alterations in their subsequent balanced CD-fed progenies.

In summary, our findings reveal that maintaining a WD for several generations promotes a progressive accumulation of epigenetic alterations in somatic and germ cells throughout generations. Two lines of evidence support this conclusion. First, ancestral exposure influences the magnitude of the overweight phenotype. Indeed, a male whose father, grandfather, great grandfather, great-great-grandfather and great-great-great grandfather, up to 5 generations of exposure, have been fed a WD exhibits the most severe overweight phenotype associated with serious metabolic alterations. Second, the father’s ancestral history (whether his ancestors were fed an unbalanced diet) affected the pattern of inheritance of this metabolic pathology.

Although it is well described that the development of type 2 diabetes is positively associated with body weight [26], we did not observe a strong correlation between fat mass and glucose and insulin sensitivities in males obtained after multigenerational WD feeding. However, we identified one obese-associated pathology that increased in severity with successive generations of a WD, namely, hepatic steatosis. Since the diet we used was not described to induce such disease, the appearance of this phenotype after multigenerational WD feeding strongly indicates that exposure sensitivity is heightened by multiple generations of exposure, at least for this diet-associated pathology. Thus, the family food environment, parental dietary behaviors and family obesity might be an additional clue to explain the increasing incidence of nonalcoholic fatty liver disease in humans [27].

Importantly, multiple generations of WD exposure impact not only the sensitivity to a WD but also the hereditary makeup, also called background. Indeed, when the father has no WD-fed ancestor, the fatness of its progenies tends to disappear after WD removal. However, in the case of fathers with several WD-fed ancestors, the progenies will remain stably overweight for more than 4 generations. Intriguingly, although the male progenies of the third and fourth generations of WD5 males were overweight, they did not develop metabolic alterations, such as glucose/insulin sensitivity alterations and fatty liver disease. Together, these data strongly suggest that the combination of ancestral and individual diet exposure was both necessary and sufficient to elicit the most severe metabolic effects in mice.

Overall, our findings are in agreement with those of recent studies of multigenerational exposure performed in several animal models. For instance, in guppies, a wide range of plastic responses under different light conditions were observed, which were dependent on multigenerational exposure to different light environments [28]. In mites, zinc element sensitivity increased by continuous multigenerational exposure [29]. In mice, male sensitivity to environmental estrogens was enhanced by successive generations of exposure [30]. Finally, rats undernourished for 50 generations showed multiple metabolic alterations that were not reversed in their respective F1 and F2 CD-fed progenies [31]. Together, the present study and previously published studies indicate that the exacerbation of stress-induced phenotypes upon multigeneration exposure as well as the stabilization of newly induced phenotypes is an evolutionarily conserved process. However, the explicit molecular mechanism(s) of this process is still largely unknown.

Single-generation exposure to a WD studies strongly indicates that sperm RNAs are a possible epigenetic vector of intergenerational epigenetic inheritance of metabolic diseases. However, these data do not exclude the possible involvement of epigenetic modifications, namely, DNA methylation, histone modifications and chromatin structure alterations. This study takes a step further in this direction, showing that sperm RNAs are vectors of intergenerational inheritance but are not sufficient for the transgenerational inheritance of diet-induced metabolic alterations. In this context, our transcriptome profile of gWAT may provide important avenues to dissect the potential molecular mechanism(s) involved in this process, revealing an enrichment in genes potentially regulated by H3K4/K27 methylation and the PRC2 complex **(Supplementary Table 2)**.

Finally, in the present study, we focused our analyses on perigonadal adipose tissue, glucose/insulin sensitivity and liver alterations. Considering the healthy and economic consequences of obesity and its comorbidities, such as cardiovascular diseases and fertility abnormalities, future studies will be important to determine the impact of multigenerational ancestor exposure on the development of obesity-associated comorbidities.

In conclusion, environmentally induced epigenetic modifications in germlines would contribute to the environmental adaptation and evolution of animal species. In the future, it will be important to assess how each epigenetic vector for inheritance interacts together to modulate the embryonic epigenome.

## Materials and Methods

### Mice

All mouse experiments were performed with C57BL/6J mice obtained from Charles River (Charles River Laboratories, France). All mice were housed in a temperature-controlled system and maintained on a 12-h light/dark cycle (lights on at 7 a.m.). Experimental mice were given *ad libitum* access to either a high-fat high-sugar diet (WD) (235 HF 45% of energy from fat, SAFE, France) or a control diet (CD) (SAFE A04, 5% of energy from fat, SAFE, France) and sterile water. To evaluate the impact of the diet of paternal ancestors on metabolic health, we developed two experimental models. On the one hand, WD feeding was maintained for 5 successive generations through the paternal line. Briefly, ten 3-week-old male mice were divided into 2 groups. Males from the first group were kept on CD, and the males of the other group were fed a WD for 3-4 months. This first generation of WD males was named WD1. At four months old, 4 to 6 independent males of each group were then crossed with 7-week-old C57BL/6J female mice (CD-fed) obtained from Charles River (Charles River Laboratories, France). The male progenies were kept and subjected to the same experimental procedure. At 3 weeks old, they were fed a WD and at 4-5 months crossed with CD-fed females. This second generation of males was called WD2. This experimental design was repeated 3 times to obtain the WD5 group **(Fig 1A)**. On the other hand, half of the WD1 and WD5 male and female progenies were fed a CD. The first generation was called F1-WD1 and F1-WD5, respectively. The F1 4-month-old male progenies were crossed with 7-week-old C57BL/6J female mice (CD-fed) to obtain the F2-WD1 and F2-WD5 progenies **(Fig 3A)**. This experimental design was repeated once to obtain the F3-WD1 and F5-WD5 progenies.

The complete experimental design was performed twice at approximately 6 months’ interval. To evaluate the role of sperm RNAs in transgenerational epigenetic inheritance of metabolic alterations, sperm RNAs extracted from 2 different CD, WD1 and WD5 males were microinjected into zygotes at the Center for Transgenic Models (University of Basel, Switzerland) following the same procedure as described in [4]. The resulting progenies were called F1-RNA-CD, F1-RNA-WD1 and F1-RNA-WD5 progenies, respectively. F2-RNA and F3- RNA progenies were obtained after crossing F1-RNA and F2-RNA 4-month-old males, respectively, with 7-week-old C57BL/6J female mice (CD-fed) obtained from Charles River (**Fig 4A)**.

All mouse experiments were conducted in accordance with the French and European legislations for the care and use of research animals.

### Body weight and food intake

Body weights were measured every week from weaning until 5 months of age. Daily food consumption was estimated by weighing the remaining food every week.

For organ measurement, 5-month-old mice were anesthetized with sodium pentobarbital and rapidly dissected. Then, gonadal WAT, inguinal subcutaneous WAT, epididymis, liver and kidneys were carefully isolated, cleaned of unrelated materials and weighed. One part was fixed in 4% PFA, and the other portion was snap frozen in liquid nitrogen.

### Blood metabolic parameter measurements

Blood metabolic parameters were detected under different physiological conditions, i.e., a random-fed state and a 16-h fasted state. Whole-blood glucose levels were determined using the OneTouch Vita (LifeScan, Johnson & Johnson company) system from tail blood. For plasma preparation, the blood was collected from the orbital sinus into sterile 1.5-ml tubes containing 2 drops of citrate sodium (3 M) and mixed gently. Blood cells were removed by centrifugation at 2000x*g* for 10 min at 4°C, and the resulting supernatant was immediately aliquoted and stored at −80°C. Serum CRP, leptin, adiponectin and cholesterol levels were measured with the C-Reactive Protein ELISA (Mouse CRP, Elabscience, CliniSciences S.A.S., Nanterre, France), Leptin ELISA (ASSAYPRO, CliniSciences S.A.S., Nanterre, France), Adiponectin ELISA (mouse Adiponectin, EZMADP-60K, EMD Millipore Corporation, Darmsbalt, Germany) and Cholesterol Assay (Abcam, Paris, France) kits, respectively. All measurements were performed in accordance with the manufacturers’ instructions.

### Glucose and insulin tolerance tests

Mice were placed in new cages prior to starvation. For GTTs, 12-h fasted mice were injected i.p. with a solution of sterile glucose (2 g/kg body weight) freshly prepared in 0.9% sterile saline. For ITTs, 6-h fasted mice were injected i.p. with insulin diluted to 0.08 mU/μl in sterile saline for a final delivery of 0.8 mU/g body weight. Baseline glucose measurements were analyzed from tail blood before i.p. glucose or insulin injection (2 mg/g body weight) using the OneTouch Vita (LifeScan, Johnson & Johnson company) system. Blood glucose measurements were taken from the tail blood at the indicated points.

### gWAT morphometry staining

gWAT was fixed with Antigenfix (Microm Microtech, France), embedded in paraffin, sectioned and stained with a hematoxylin and eosin solution. Slides (4/group) were scanned with Axio- scan, which allowed the scanning of the entire slide at high resolution. Six pictures of six different areas from 1–2 sections per sample were chosen and analyzed with image analyzer software (ImageJ). Total areas of adipocytes were traced manually. The total count ranged from 3275 to 7052 adipocytes per condition. The mean surface area of the adipocytes was calculated using image analyzer software (ImageJ). For each sample, 400–1000 adipocytes were counted.

### Estimation of adipocyte number in gWAT

To estimate the number of adipocytes in gWAT depots, we applied a mathematical equation developed by Jo and colleagues[32], as previously described in [20]. Briefly, the number of adipocytes (N) was estimated by dividing the WAT mass (M) by the density of adipocytes (D = 915 g/L) multiplied by the mean volume of adipocytes within the WAT (V). The mean volume of adipocytes is calculated from the mean diameters of adipocytes, extracted from tissue sections images. The equation is presented below:

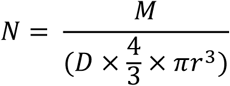

### Computed tomography of mice

Anesthetized animals were placed in a SkyScan μCT-1178 X-ray tomograph (Bruker) and analyzed as previously described[33]. Mice were scanned using the following parameters: 104 µm pixel size, 49 kV, 0.5-mm-thick aluminum filter and a rotation step of 0.9°. 3D reconstructions and analysis of whole abdominal fat were performed using NRecon and CTAn software (Skyscan), respectively, between thoracic 13 and sacral 4 vertebral markers.

### Liver triglyceride Measurement

Frozen small piece of liver was placed in 2ml tubes with Ceramic Beads (for Precellys homogenizer) and were homogenized in Sodium Acetate (0.2M, pH4.5) using the Precellys homogenizer. After centrifugation, the supernatant was stored at −80°C. The TG in homogenates was measured according to the reagent kit instruction (Triglycerides FS - DiaSys Diagnostic Systems GmbH, Holzheim, Germany).

### Histological liver examination

The livers were prepared and fixed in 4% paraformaldehyde, embedded in paraffin, cut into 5- μm-thick slices, stained with haematoxylin and eosin (H&E), mounted with neutral resins and then scanned with Axio-scan, which allowed the scanning of the entire slide at high resolution. Liver histology was blindly evaluated by two independent analysts using a semiquantitative scale adapted from previously validated procedures [34]. To that end, images from three different fields in each section were collected at 20× magnification and numbers of normal hepatocytes, microvesicular and macrovesicular steatosis and degenerating hepatocytes was assessed.

### Sperm collection

Sperm were collected from the epididymis by squeezing. The cell suspension was centrifuged at 1000 rpm for 5 min, and the supernatant containing the spermatozoa was centrifuged at 3000 rpm for 15 min. To reduce contamination of somatic cells, the pellet was submitted to hypotonic shock by resuspension in water (250 μl), followed by the addition of 15 ml of PBS. The suspension was finally centrifuged at 3000 rpm for 15 min.

### Quantitative RT-PCR

Total RNA from epididymal adipose tissues was extracted using TRIzol reagent (Life Technologies, France) according to the manufacturer’s instructions. Total RNA (0.5 µg) was reverse transcribed with mouse myeloblastosis virus reverse transcriptase (Promega) under standard conditions using hexanucleotide random primers according to the manufacturer’s instructions. cDNA was amplified by PCR with specific primers. Real-time PCR was performed on the Light Cycler Instrument (Roche Diagnostics) using the Platinum SYBR Green kit (Invitrogen). Specific primers for mouse leptin and 2 mouse housekeeping genes used for normalization (*β-actin* and *34B4* mouse genes) were purchased from Sigma (Sigma, France). We used primers for *Leptin (forward*, AAC CTG GAA ATG CTC TGG CTGT; *reverse*, ACT CGC TGT GAA TGG CCT GAA A), *36B4F (forward*, TCC AGG CTT TGG GCA TCA; *reverse*, CTT TAT CAG CTG CAC ATC ACT CAG A), and *β-actin* (forward, CTA AGG CCA ACC GTG AAA AG; reverse, CCT GCT TCA CCA CCT TCT TG).

### RNA preparation and microinjection

Frozen sperm were stored at −80°C. RNA was then extracted by the TRIzol procedure (Invitrogen). The same preparations of sperm RNAs were used for microinjection and small RNA sequencing. RNA preparations were verified by spectrometry on an Agilent Bioanalyzer 2100 apparatus. Microinjection into fertilized eggs was performed as described in [25]. RNA solutions were adjusted to a concentration of 1-2 µg/ml, and 1-2 pl were microinjected into the pronucleus of C57BL/6 fertilized mouse oocytes.

### Library preparation and sequencing

Total RNA was isolated from gonadal adipose tissue (eWAT; n = 9) samples using the Ambion RiboPure (Thermo Fisher Scientific). RNA was quantified in a Nanodrop ND-1000 spectrophotometer and RNA purity and integrity was checked by using a Bioanalyzer-2100 equipment (Agilent Technologies, INC., Santa Clara, CA). Libraries were prepared using the TruSeq RNA Sample Preparation Kit (Ilumina Inc., CA) and were paired-end sequenced (2 × 75 bp), by using the TruSeq SBS Kit v3-HS (Illumina Inc., CA), in a HiSeq 2000 platform (Illumina Inc., CA). More than 30 M PE reads were obtained for all samples.

### Transcriptomics analysis (RNA-sequencing analysis)

Raw sequence files were subjected to quality control analysis using FastQC. In order to avoid low quality data, adapters were removed by Cutadapt and lower quality bases were trimmed by trimmomatic [35]. The quality-checked reads processed were mapped to the mouse reference genome GRCm38/mm10 using STAR [36]. Reads abundance was evaluated for each gene followed by annotation versus mouse GTF by using the feature Counts function. The R package Edger was used in order to normalize the reads and to identify differentially expressed (DE) genes [37]. Genes with FDR < 0.05 after correcting for multiple testing were classified as DE [38]. The pheatmap and VolcanoPlot functions (R packages) were used to graphically represent the expression levels (log2FC) and significance of DE genes among treatments. These experiments have been deposited in the GEO Database with accession number (GSE148972) and a review access token (ovwzywcgnpublor).

### Small RNA-sequencing analysis: Analysis of differential expression

The experiment was carried out in triplicate. RNA libraries were prepared starting from 50-100 ng of total RNA from individual mice (n=3 per group, 3 groups in total) and constructed using the Illumina TruSeq Stranded Small RNA Sequencing kit (Illumina) according to the manufacturer’s instructions. Sequencing was performed at the IPMC platform (Sophia-Antipolis, France) using the HiSeq 2500 (Illumina).

Read quality was assessed using FastQC and trimmed, against known common Illumina adapter/primer sequences, using trimmomatic. The SmallRNAs IPMC pipeline with Illumina adaptor trimming was used, read sizes < 15 b were discarded. Reads kept were mapped to the mouse genome GRCm38/mm10 by using bowtie2 (--local --very-sensitive-local -k 24). Reads abundance was evaluated for each gene followed by annotation versus gff mirbase v21, ensembl ncrna rel73, tRNAs and piRNA clusters from piRNAclusterDB. Normalization of reads abundance and differential expression analysis was performed by using DESeq R package. The baseMean for each gene, the maximum of mean counts among all conditions, was at least 50 counts. NGS experiments have been deposited in the GEO Database with accession number (GSE138989).

### Statistics and reproducibility

Statistical analyses were performed using the Kruskal-Wallis test followed by the two-stage step-up method of Benjamini, Krieger and Yekuteil for multiple comparisons of body weight, body composition, cholesterol, and leptin levels, as well as leptin mRNA expression and AUC- GTT and AUC-ITT between the WD cohorts, F1-, F2-, and F3-progenies and RNA- microinjected progenies.

To measure the linear relationship between two variables, we used Spearman’s correlation coefficient. All statistical analyses were performed with Prism 7 for Mac OS X software (GraphPad software, Inc.). Data are presented as the median ± SD. A *p* value of <0.05 was considered statistically significant.

Sample size and replicates are indicated in the figure legends. The WD cohort and WD progenies were repeated twice.

## Acknowledgments

We are grateful to Dr Jean-Jacques Remy for his careful help from the start of this project. We thank Dr Mireille Cormont, Sofia Fazio, Dr Maria Stathopoulou and Dr Claire Mauduit for constructive discussions. We thank Marion Dussot for her technical assistance in performing the liver biochemistry. We relied on sequencing data generated by the IPMC Functional Genomics Facility (UCAGenomiX - IPMC platform (Sophia-Antipolis, France)). We thank the Center for Transgenic Models (University of Basel, Switzerland) for the mouse microinjection assays. We are grateful to the C3M mouse facility (U1065, Nice). This work has been supported by ANR (grant# ANR-12-ADAPT-0022) and the FFAS “Fonds Français pour l’Alimentation et la Santé “(15D52) and was partly supported by research funding from the Canceropôle PACA, Institut National du Cancer and Région Sud. F.S. was supported by the UCA-IDEX.

## Author Contributions

V.G. conceived and designed the project. V.G., G.R., J.G., V.C.M., D.P., E.Z.A. and M.A.D. performed the experiments. V.G. wrote the manuscript. F.S., G.R., J.G., E.Z.A and F.S. contributed to the data analysis. E.Z.A., D.P.V., C.M., F.S., G.R., L.M. and M.T. edited the paper. All of the authors read and approved the final manuscript.

## Figure Legends

**S1 Fig. Exacerbation of the overweight phenotype upon continuous paternal WD feeding for multiple generations.**

(A) Evolution of total body weight at 12 and 18 weeks over WD-fed generations (n≥15). (B) Positive linear correlation between gWAT and total body weight. Statistically significant positive linear correlation between gWAT and plasma leptin concentration (C), between gWAT and plasma total cholesterol (D), between gWAT and plasma CRP concentration (E) and between plasma leptin concentration and gWAT leptin mRNA (F). (G) There was no statistically significant linear correlation between epididymal fat mass and AUC-GTT. (H) Principal component analysis (PCA) analysis of fasting blood glucose, total body weight, epididymal fat mass and kCal in the different WD cohorts visualizing the pattern of WD males depending on the number of WD-fed ancestors.

**S2 Fig. Long-term epigenetic inheritance of a “healthy” overweight phenotype in CD-fed progenies from WD5 males.**

Box-whiskers (min-max) of the median total body weight of the male (A) and female (C) F1-, F2-, F3-WD progenies (n≥8 mice per group). Box-whiskers (min-max) of the median gonadal fat mass (gWAT) weight relative to total body weight in the male (B) and female (D) F1-, F2-, F3-WD progenies (n≥8 mice per group). The evolution of glucose parameters in CD-fed male (E, F) and female (G, H) WD progenies. Above the glucose tolerance curves are representative corresponding box-whiskers (min-max) of the median AUC-GTT of each group (E, G). Above the insulin tolerance curves are representative corresponding box-whiskers (min-max) of the median AUC-ITT of each cohort (F, H) measured in each WD cohort.

*p<0.05, **p<0.01, ***p<0.001 (The Kruskal-Wallis test, a rank-based nonparametric test for multiple comparisons, two-stage linear step-up procedure of Benjamin, Krieger and Yekutieli was used to calculate the adjusted *p* value). § denotes the WD5 progeny whose median was significantly different from that of the WD1 corresponding progeny.

**S3 Fig. No liver alteration was observed in the CD-fed progenies from WD1 and WD5 males.**

(A) Liver triglyceride contents in CD, WD1 and WD5 CD-fed progenies. (B) H&E staining of liver sections (scale bar: 250 μm) in representative F1, F2, and F3 CD-fed progenies of CD, WD1 and WD5 males.

**S4 Fig. Small RNA-seq analysis of WD1 and WD5 male sperm**

(A) Representative bioanalyzer profiles of CD, WD1 and WD5 sperm total RNAs. (B) The normalized small RNA levels from the CD (blue spots), WD1 (red spots) and WD5 (green spots) sperm were analyzed by PCA. One WD5 fell outside the PCA cluster and was arbitrarily removed for differential expression analysis. (C) Venn diagram of small RNA sequences differentially expressed in WD1 and WD5 sperm. The numbers of small RNAs that are unique for each WD1 and WD5 male are shown in each circle. The numbers of genes in overlapping (common) are indicated at the intersections of the sets in the Venn diagram (P_adjvalue_<0.05 Log2FC≥|0.6|). Heatmap diagrams of microRNAs (D) or piRNAs, tRNA fragments, and other small RNAs (E) differentially expressed (P_adjvalue_<0.05 Log2FC≥|0.6|) in both WD1 and WD5 sperm compared to their expression in the CD sperm cohort. The color green indicates a negative deregulation, whereas red shows a positive deregulation.

CD, chow diet; WD1, male fed a WD for one generation; WD5, male fed a WD for 5 successive generations.

**S5 Fig. No liver alteration was observed in the CD-fed progenies from RNA-WD1 and RNA-WD5 males.**

(A) Liver triglyceride contents in F1, F2, and F3 progenies of CD-RNA-, WD1-RNA- and WD5- RNA-injected groups. (B) H&E staining of liver sections (scale bar: 250 μm) in representative F1, F2 and F3 CD-fed progenies of CD-RNA-, WD1-RNA- and WD5-RNA-injected groups.

