## Supplementary information for "Paternal multigenerational exposure to an obesogenic diet drives epigenetic predisposition to metabolic disorders"

**S1 Table Physiological characteristics of different WD groups**

| Characteristic | Control n=17 | WD1 n=15 | WD2 n=15 | WD3 n=16 | WD4 n=42 | WD5 n=53 |
|---|---|---|---|---|---|---|
| Body weight (g) (12 weeks) | 25.7±1.0 | 25.9±1.4 | 26.7±1.8 | 26.6±1.8 | **28.2±2.2***§** | **27.6±1.8*** §** |
| Body weight (g) (16 weeks) | 27.7±0.2 | **29±1.4*** | **29.9±2.6*** | 28.5±1.7 | **31.1±2.7***§** | **30.8±2.5***§** |
| Daily diet consumption (Kcal/day) | 10.3±0.3 | **11.6±1.8***** | **12.1±1.3***** | **11.6±0.6***** | **12.1 ±1.4***** | **11.4±0.8***** |
| Kidney (g) | 0.37±0.03 | 0.4±0.08 | 0.4±0.06 | 0.4±0.06 | **0.43±0.09**** | **0.42±0.08*** |
| Kidney/body weight (%) | 1.3±0.15 | 1.4±0.23 | 1.39±0.19 | 1.42±0.15 | 1.37±0.31 | **1.44±0.33*** |
| Heart (g) | 0.2±0.05 | 0.2±0.04 | 0.18±0.04 | 0.18±0.07 | 0.21±0.03 | 0.21±0.2 |
| Liver (g) | 1.38±0.19 | 1.49±0.27 | 1.53±0.22 | 1.40±0.38 | **1.86±0.40***§** | **1.7±0.34**** |
| Liver/body weight (%) | 5±0.49 | 5.2±0.66 | 5.3±0.59 | 5.4±0.48 | **5.8±0. 76***§** | **5.6±0. 79**** |
| gWAT (g) | 0.41±0.12 | **0.92±0.45**** | **0.97±0.28***** | **0.97±0.97***** | **1.25±0.52***** | **1.24±0.32***§** |
| gWAT/body weight (%) | 1.3±0.35 | **3.18±1.3**** | **3.3±0.8***** | **3.3±0.8***** | **4.0±1.34***** | **4.1±1.06***** |
| Abdominal Adipose Volume (arbitrary unit) | 1465±612 (n=10) | 2660±496 (n=5) | **2471±268* (n=4)** | **3499±1007** (n=6)** | **3694±1173** (n=7)** | **4311±1498*** (n=11)** |

Values are expressed as mean ±SD. Kruskal-Wallis test, a rank-based non-parametric test for multiple comparisons, two-stage linear step-up procedure of Benjamin, Krieger and Yekutieli was used to calculate the *p* value. Numbers are in bold if p<0.05. * and § identified the WDs groups whose mean rank difference was statistically significantly different as compared to that of the CD and WD1 groups, respectively. *p<0.05, **p<0.01, ***p<0.001.

**S3 Table Physiological characteristics of F1, F2 and F3 male progenies from either WD1 or WD5 males**

| Characteristic | Control n=20 | F1 | | F2 | | F3 | |
|---|---|---|---|---|---|---|---|
| | | WD1 n=10 | WD5 n=17 | WD1 n=14 | WD5 n=17 | WD1 n=10 | WD5 n=17 |
| Body weight (g) (12 weeks) | 25.6±1.01 | **28.3±1.01*** ** | **29.3±2.6*** ** | 27.15±1.01 | **28.9±1.3***§** | 25.4±1.0 | **29.4±1.8**§** |
| Body weight (g) (18 weeks) | 27.2±4 | **30.2±1.4*** | **29.4± 1.7*** | **30.5±1.8**** | **31.4±1.4***** | 27±1.8 | **31.5±2.6***§** |
| Food intake (Kcal/d) | 10.3±0.3 | **11.3±0.2**** | 10.1±0.2 | 10.6±0.2 | **11.5±1.0***** | 10.3±0.3 | 9.8±1.0 |
| Kidney (g) | 0.37±0.03 | 0.36±0.04 | 0.39±0.05 | 0.38±0.05 | 0.37±.0.03 | 0.37±0.03 | **0.45±0.06*§** |
| Kidney/body weight (%) | 1.3±0.15 | 1.2±0.08 | 1.3±0.13 | 1.4±0.16 | 1.2±0.11 | 1.3±0.11 | **1.1±0.1*** |
| Heart (g) | 0.2±0.05 | 0.2±0.03 | 0.22±0.04 | 0.2±0.02 | 0.20±0.03 | 0.2±0.01 | 0.20±0.03 |
| gWAT (g) | 0.43±0.12 | **0.48±0.05*** | **0.61±0.26*** | **0.72±0.17*** | **0.66±0.2***** | 0.34±0.08 | **0.82±0.2***§** |
| gWAT/body (%) | 1.25±0.35 | **1.6±0.20*** | **1.7±0.44*** | **2.3±0.9***** | **2.1±0.6***** | 1.2±0.19 | **3.4±1.4***§** |
| Liver (g) | 1.43±0.02 | 1.47±0.19 | **1.49±0.20*** | 1.41±0.26 | **1.6±0.19*** | 1.44±0.02 | **1.7±0.22*** |
| Liver/body weight (%) | 5.06±0.1 | 4.8±0.1 | 4.65.06±0.2 | **4.5±0.1*** | 4.95.06±0.1 | 5.06±0.1 | 4.8±0.4 |
| Fasting Glucose (g) | 83±10 | 71±13 | 79±13 | **69±11*** | **68±6.7*** | 76±3 | 82±21 |
| AUC-GTT (mg/dl/min) | 21886±3738 | **24756±3234** | **25977±3445*** | **25756±4957** | 23238±4258 | 20203±24812 | 21861±5616 |
| AUC-ITT (mg/dl/min) | 3136±271 | 3702±475 | 3106±354 | 4230±1242 | 3041±736 | 3200±271 | 3327±914 |
| Total cholesterol (mg/dl) | 1.06±0.35 | 0.9±0.20 | 0.91±0.3 | 0.67±0.2 | 0.84±0.12 | nd | nd |
| Abdominal Adipose Volume (mm3) | 1463±612 n=9 | **1732±321* n=6** | **2122±573** n=7** | nd | **2738±1127** n=5** | nd | nd |

Values are expressed as mean ± SD. Kruskal-Wallis test, a rank-based non-parametric test for multiple comparisons, two-stage linear step-up procedure of Benjamin, Krieger and Yekutieli was used to calculate the *p* value. Numbers are in bold if p<0.05. * and § identified the WDs groups whose mean rank difference was statistically significantly different as compared to that of the CD and WD1 groups, respectively. *p<0.05, **p<0.01, ***p<0.001, nd= not determined.

**S4 Table Physiological characteristics of F1, F2 and F3 female progenies from either WD1 or WD5 males**

| Characteristic | Control n=11 | F1 | | F2 | | F3 | |
|---|---|---|---|---|---|---|---|
| | | WD1 n=10 | WD5 n=15 | WD1 n=9 | WD5 n=15 | WD1 n=7 | WD5 n=13 |
| Body weight (g) (12 weeks) | 20.2±1.5 | **22.7±1.2*** | 22.2±0.3 | 20.4±1.2 | **21.6±1.1*§** | 21.0±0.4 | **22.9±1.4***§** |
| Body weight (g) (18 weeks) | 21.4±0.8 | **22.7±1.5*** | 22.2±1.7 | 22.4±1.5 | **23.5±1.8**** | 21.4±0.4 | **24.08±1.6**§** |
| Food intake (Kcal/d) | 8.8±0.2 | 9.2±0.1 | 9.3±0.2 | 9.5±0.1 | 9.3±0.2 | 8.6±0.3 | **9.5±0.2***§** |
| Kidney (g) | 0.26±0.02 | 0.27±0.03 | 0.29±0.05 | 0.3±0.06 | 0.27±003 | 0.26±0.01 | 0.28±0.02 |
| Kidney/body weigt (%) | 1.22±0.07 | 1.2±0.1 | 1.3±0.18 | 1.27±0.05 | 1.1±0.12 | 1.1±0.02 | 1.2±0.23 |
| Heart (g) | 0.2±0.05 | 0.2±0.03 | 0.2±0.04 | 0.2±0.04 | 0.20±0.03 | 0.2±0.02 | 0.2±0.03 |
| gWAT (g) | 0.28±0.44 | 0.36±0.11 | **0.37±0.37*** | 0.26±0.09 | **0.65±0.3**§** | 0.27±0.03 | **0.44±0.29***§** |
| gWAT/body weight (%) | 1.09±0.09 | **1.57±0.35*** | 1.66±0.55 | 1.28±0.19 | **2.3±1.3*** | 1.1±0.02 | **1.7±0.6*§** |
| Liver (g) | 1.1±0.08 | 1.1±0.11 | 0.9±0.21* | 1.1±0.27 | 1.1±0.08 | 1.1±0.02 | 1.1±0.07 |
| Liver/body weight (%) | 5.08±0.5 | 4.5±0.5 | **3.9±0.67*** | **4.5±0.61*** | 4.7±0.5 | 4.8±0.5 | 4.8±1.05 |
| Fasting Glucose (g) | 67±11 | 82±13 | 85±22 | 68±10 | 71±9 | 70±7 | 69±15 |
| AUC-GTT (mg/dl/min) | 15614±2210 | 18897±3390 | **21617±3354*** | **27984±8669***** | 18271±2585§ | 18241±3864 | 21404±3864 |
| AUC-ITT (mg/dl/min) | 3136±187 | 3702±280 | 3106±354 | **4230±450*** | 3041±256 | 3200±129 | 3291±761 |
| Total Cholesterol (mg/dl) | 0.57±0.09 | 0.63±0.15 | 0.59±0.17 | nd | 0.69±0.05 | nd | nd |

Values are expressed as mean ± SD. Kruskal-Wallis test, a rank-based non-parametric test for multiple comparisons, two-stage linear step-up procedure of Benjamin, Krieger and Yekutieli was used to calculate the *p* value. Numbers are in bold if p<0.05. Numbers are in bold if p<0.05. * and § identified the WDs groups whose mean rank difference was statistically significantly different as compared to that of the CD and WD1 groups, respectively. *p<0.05, **p<0.01, ***p<0.001, nd= not determined.

**S7 Table Physiological characteristics of F1 male and female progenies RNA microinjected embryos**

| Characteristic | F1-RNA male progenies | | | F1-RNA female progenies | | |
|---|---|---|---|---|---|---|
| | **RNA-CD n=9** | **RNA-WD1 n=12** | **RNA-WD5 n=38** | **RNA-CD n=10** | **RNA-HFD1 n=6** | **RNA-WD5 n=30** |
| Body weight (g) (10 weeks) | 27.55±1.2 | 28.45±1.8 | **29.88±2.08*****| 22.58±0.9 | 22±1.37 | 22±1.16 |
| Body weight (g) (12 weeks) | 30.18±1.05 | **31.36±1.83*** | **31.05±2.07*** | 23.5±0.9 | 23.2±1.6 | 23.4±1.5 |
| Body weight (g) (16 weeks) | 33.8±0.9 | 33.4±1.9 | 33.1±2.7 | 26.7±1.3 | 25.5±2.5 | 25.6±2.6 |
| Kidney (g) | 0.5±0.01 | 0.46±0.08 | 0.49±0.1 | 0.31±0.06 | 0.3±0.03 | 0.32±0.08 |
| Kidney to body mass ratio (%) | 1.35±0.08 | 1.25±0.21 | 1.41±0.37 | 1.06±0.07 | 1.09±0.03 | 1.23±0.04 |
| gWAT (g) | 0.95±0.47 | 0.85±0.36 | 1.14±0.57 | 0.74±0.24 | 0.62±0.20 | 0.78±0.43 |
| gWAT to body mass ratio (%) | 2.52±0.9 | 2.47±0.8 | 2.49±0.8 | 2.4±0.9 | 2.3±0.8 | 2.6±1.1 |
| Liver (g) | 1.57±0.38 | 1.7±0.37 | 1.8±0.3 | 1.4±0.24 | **1.1±0.05**** | 1.4±0.16 |
| Liver to body mass ratio (%) | 4.2±0.8 | 4.8±0.8 | 4.2±0.7 | 4.8±0.7 | **4.1±0.4*** | 4.7±0.4 |
| Fasting Glucose (mg/dl) | 104±15.6 | 104±8.5 | **128±17*** | 112±15 | **121±21*** | 99±15 |
| AUC-GTT (mg/dl/min) | 33867±546 | **36128±1434** | **37002±939*** | 29215±1151 | **32428±1482*** | 31767±1244 |
| AUC-ITT (mg/dl/min) | 6468±976 | **8751±490** | **8139±349** | 7677±131 | 7028±409 | 7855±315 |

Values are expressed as mean ± SD. Kruskal-Wallis test, a rank-based non-parametric test for multiple comparisons, two-stage linear step-up procedure of Benjamin, Krieger and Yekutieli was used to calculate the *p* value. Numbers are in bold if p<0.05. Numbers are in bold if p<0.05. * identified the WDs groups whose mean rank difference was statistically significantly different as compared to that of the CD group. *p<0.05, **p<0.01, ***p<0.001.

**S8 Table Physiological characteristics of F2 male and female progenies RNA microinjected embryos**

| Characteristic | F2-RNA male progenies | | | F2-RNA female progenies | | |
|---|---|---|---|---|---|---|
| | **RNA-CD n=11** | **RNA-WD1 n=11** | **RNA-WD5 n=24** | **RNA-CD n=11** | **RNA-WD1 n=8** | **RNA-WD5 n=18** |
| Body weight (g) (10 weeks) | 25.65±1.2 | **27.97±1.6*** ** | **27.17±1.26*** ** | 19.05±0.9 | 20.09±0.3 | 19.95±1.23 |
| Body weight (g) (12 weeks) | 26.95±1.23 | **29.89±1.97** ** | **28.72±1.12** ** | 22.2±0.9 | **24.09±2.1*** | 21.37±1.3 |
| Body weight (g) (16 weeks) | 29.2 ±1.38 | **31.85±3.13 *** | **31.66±1.43 ** ** | 22.98±0.9 | **25.51±2.8*** | 22.78±1.4 |
| Kidney (g) | 0.39±0.05 | 0.37±0.07 | 0.42±0.08 | 0.32±0.03 | 0.29±0.02 | 0.3±0.03 |
| Kidney to body mass ratio (%) | 1.2±0.14 | 1.04±0.15 | 1.21±0.13 | 1.14±0.05 | 1.13±0.04 | 1.01±0.08 |
| gWAT (g) | 0.54±0.27 | **0.96±0.44*** | 0.73±0.27 | 0.45±0.21 | **0.8±0.29*** | **0.9±0.4*** |
| gWAT to body mass ratio (%) | 2.17±0.6 | 3.03±1.19 | 2.49±0.7 | 1.6±0.6 | **3.6±1.0** | **3.2±1.3*** |
| Liver (g) | 1.4±0.24 | **1.9±1.8** ** | 1.6±0.20 | 1.3±0.23 | 1.3±0.22 | 1.2±0.25 |
| Liver to body mass ratio (%) | 4.7±0.65 | 5.3±0.43 | 4.8±0.7 | 4.05±0.6 | 4.9±0.6 | 4.37±0.63 |
| Fasting Glucose (mg/dl) | 184±17 | 203±14 | 156±33 | 129±14 | **148**±41 | 123±31 |
| AUC-GTT (mg/dl/min) | 31994±863 | 34880±1596 | 33076±1045 | 29350±519 | 31290±1067 | 31042±1173 |
| AUC-ITT (mg/dl/min) | 8139±349 | **12953±1926*** | 8187±795 | 5001±192 | **6172±448*** | **6607±322*** |

Values are expressed as mean ± SD. Kruskal-Wallis test, a rank-based non-parametric test for multiple comparisons, two-stage linear step-up procedure of Benjamin, Krieger and Yekutieli was used to calculate the *p* value. Numbers are in bold if p<0.05. Numbers are in bold if p<0.05. * identified the WDs groups whose mean rank difference was statistically significantly different as compared to that of the CD. *p<0.05, **p<0.01, ***p<0.001.

**S9 Table Physiological characteristics of F3 male and female progenies RNA microinjected embryos**

| Characteristic | F3-RNA male progenies | | | F3-RNA female progenies | | |
|---|---|---|---|---|---|---|
| | **RNA-CD n=6** | **RNA-WD1 n=8** | **RNA-WD5 n=15** | **RNA-CD n=12** | **RNA-WD1 n=10** | **RNA-WD5 n=13** |
| Body weight (g) (10 weeks) | 26.1±1.3 | 25.8±1.7 | 25.87±1.6 | 20.2±1.16 | 19.3±0.9 | 20.17±1.1 |
| Body weight (g) (12 weeks) | 26.9±1.35 | 27.86±1.31 | 27.65±1.39 | 20.2±1.2 | 20.87±1.5 | 21.22±1.5 |
| Body weight (g) (16 weeks) | 29.2±1.38 | 29.6±1.5 | 29.93.4±2.0 | 22.08±0.53 | 22.54±0.6 | 22.87±0.5 |
| Kidney (g) | 0.38±0.016 | 0.45±0.09 | 0.39±0.03 | 0.3±0.02 | 0.29±0.06 | 0.31±0.04 |
| Kidney (%) | 1.2±0.23 | 1.2±0.23 | 1.1±0.11 | 1.2±0.06 | 1.14±0.03 | 1.18±0.05 |
| gWAT (g) | 0.51±0.27 | 0.63±0.13 | 0.75±0.20 | 0.45±0.13 | 0.35±0.13 | 0.43±0.21 |
| gWAT (%) | 1.7±0.48 | 2.03±0.37 | 2.0±0.51 | 1.6±0.3 | 1.46±0.7 | 1.7±0.8 |
| Liver (g) | 1.4±0.26 | **1.7±0.2*** | **1.6±0.22*** | 1.3±0.1 | 1.16±0.12 | 1.26±0.27 |
| Liver (%) | 4.6±0.6 | 4.6±0.16 | 4.4±0.3 | 4.2±0.4 | 4.7±0.41 | 4.7±0.77 |
| Fasting Glucose (mg/dl) | 168±45 | 150±41 | **207±41*** | 150±29 | 174±29 | 169±28 |
| AUC-GTT (mg/dl/min) | 31493±852 | 30615±2087 | **39418±2801**** | 29350±1067 | 33614±3503 | 33553±2066 |
| AUC-ITT (mg/dl/min) | 8640±1042 | 8637±1085 | **12917±923*** | 7817±192 | **8248±857**** | **9602±7578**** |

Values are expressed as mean ± SD. Kruskal-Wallis test, a rank-based non-parametric test for multiple comparisons, two-stage linear step-up procedure of Benjamin, Krieger and Yekutieli was used to calculate the *p* value. Numbers are in bold if p<0.05. * identified the WDs groups whose mean rank difference was statistically significantly different as compared to that of the CD. *p<0.05, **p<0.01, ***p<0.001, nd= not determined.

**S10 Table Physiological characteristics of F4 male and female progenies RNA microinjected embryos**

| Characteristic | F4-RNA male progenies | | | F4-RNA female progenies | | |
|---|---|---|---|---|---|---|
| | RNA-CD n=6 | RNA-WD1 n=8 | RNA-WD5 n=10 | RNA-CD n=6 | RNA-WD1 n=6 | RNA-WD5 n=9 |
| Body weight (g) (10 weeks) | 26.9±0.6 | 26.01±2.3 | 26.81±2.3 | 20.18±1.2 | 19.15±1.0 | 20.97±1.4 |
| Body weight (g) (12 weeks) | 28.68±0.8 | 26.51±3 | 2861±1.1 | 21.8±0.9 | 20.02±1.3 | 21.34±1.1 |
| Body weight (g) (16 weeks) | 29±0.6 | 28.88±2.6 | 29.12±1.6 | 21.81±1.0 | 21.43±1.4 | 21.91±1.1 |
| Fasting Glucose (mg/dl) | 179±11 | 158±8 | 186±20 | 152±5.0 | 176±12 | 170±14 |
| AUC GTT(mg/dl) | 31060±9395 | 29645±3257 | 31033±6265 | 29350±351 | 33636±1918 | 35506±1684 |
| AUC ITT (mg/dl) | 8647±1004 | 10148±687 | 8440±1405 | 7865±554 | 8005±532 | 7653±466 |

Values are expressed as mean ± SD. Kruskal-Wallis test, a rank-based non-parametric test for multiple comparisons, two-stage linear step-up procedure of Benjamin, Krieger and Yekutieli was used to calculate the *p* value.

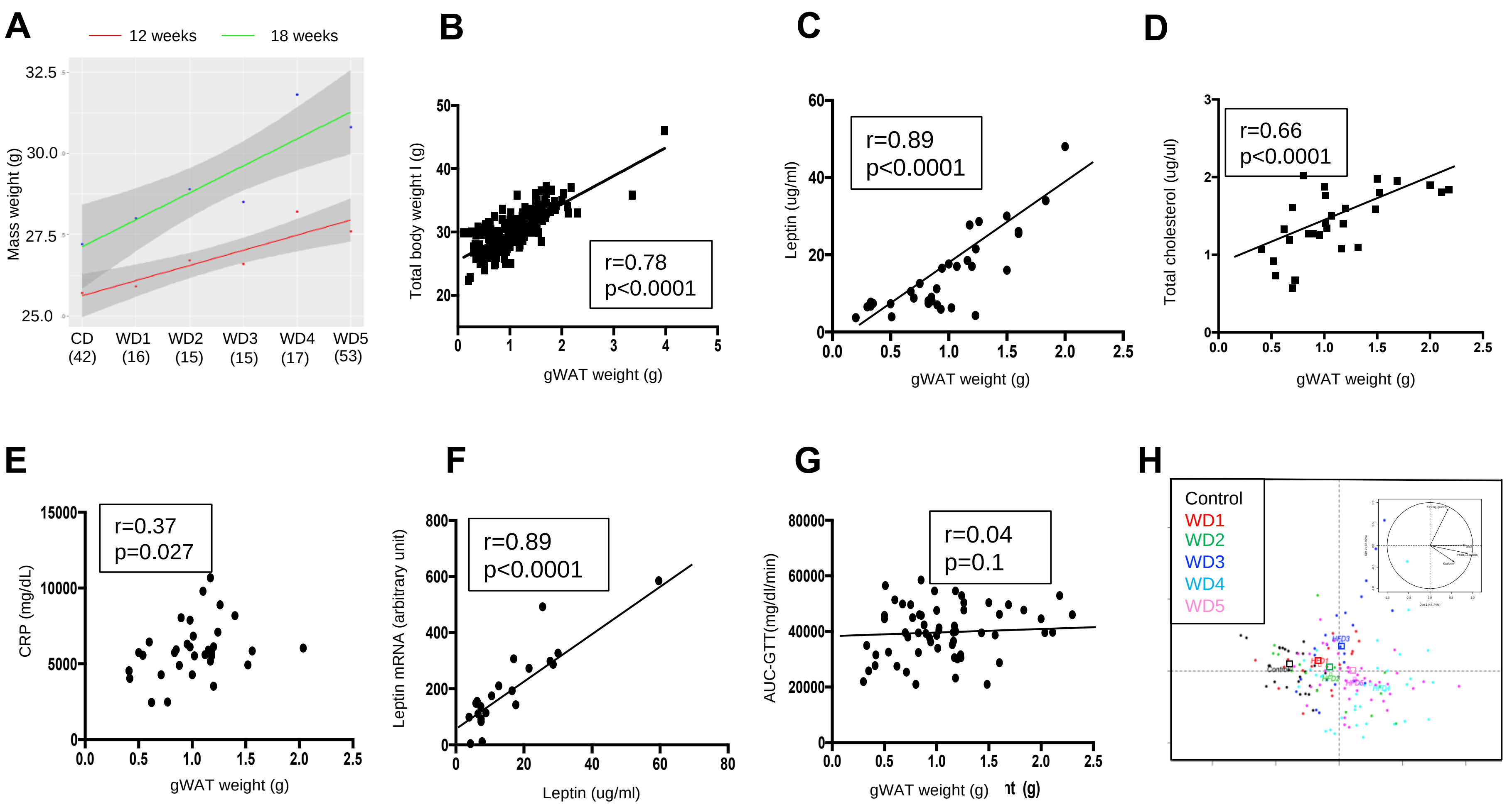
A
12 weeks
18 weeks
Mass weight (g)
CD (42)
WD1 (16)
WD2 (15)
WD3 (15)
WD4 (17)
WD5 (53)
B
Total body weight I (g)
gWAT weight (g)
r=0.78
p<0.0001
C
Leptin (ug/ml)
gWAT weight (g)
r=0.89
p<0.0001
D
Total cholesterol (ug/ul)
gWAT weight (g)
r=0.66
p<0.0001
E
CRP (mg/dL)
gWAT weight (g)
r=0.37
p=0.027
F
Leptin mRNA (arbitrary unit)
Leptin (ug/ml)
r=0.89
p<0.0001
G
AUC-GTT(mg/dl/min)
gWAT weight (g)
r=0.04
p=0.1
H
Control
WD1
WD2
WD3
WD4
WD5

S1 Fig

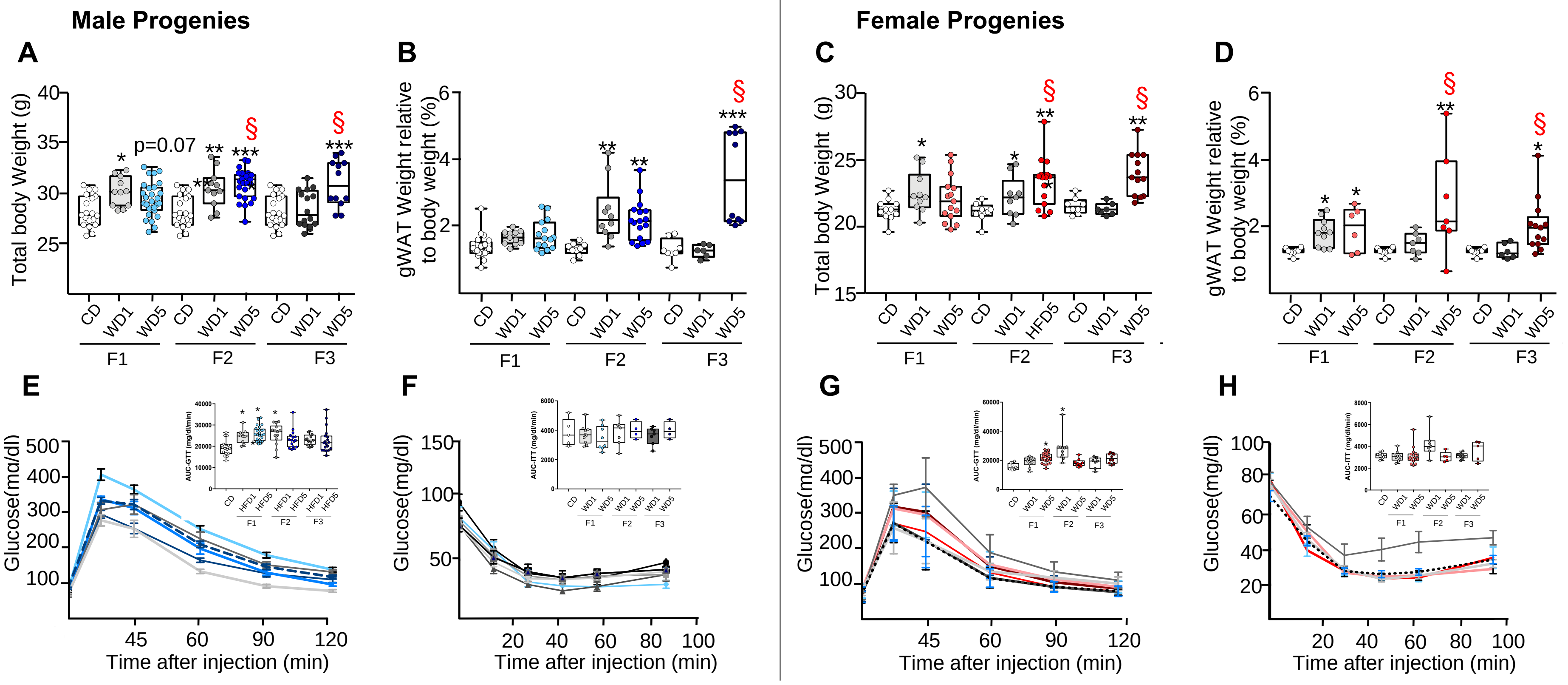
Male Progenies
Female Progenies
A
B
C
D
E
F
G
H
Total body Weight (g)
gWAT Weight relative to body weight (%)
p=0.07
CD
WD1
WD5
HFD5
F1
F2
F3
Glucose(mg/dl)
Time after injection (min)
AUC-GTT (mg/dl/min)
AUC-ITT (mg/dl/min)
HFD1

S2 Fig

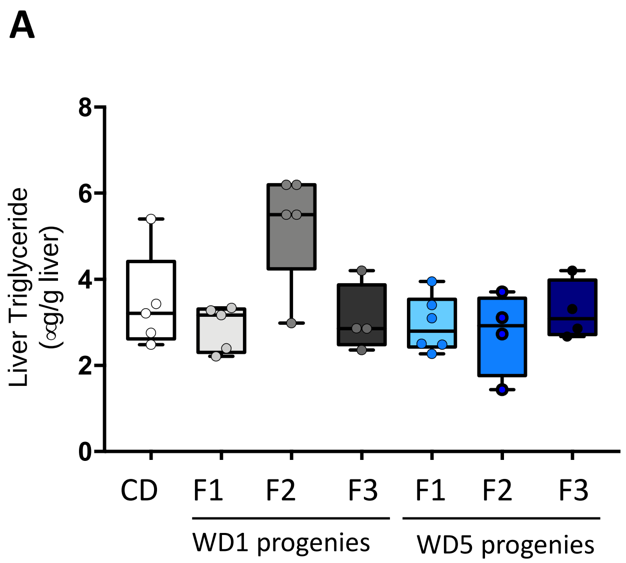
A
Liver Triglyceride (∝g/g liver)
8
6
4
2
0
CD
F1
F2
F3
F1
F2
F3
WD1 progenies
WD5 progenies

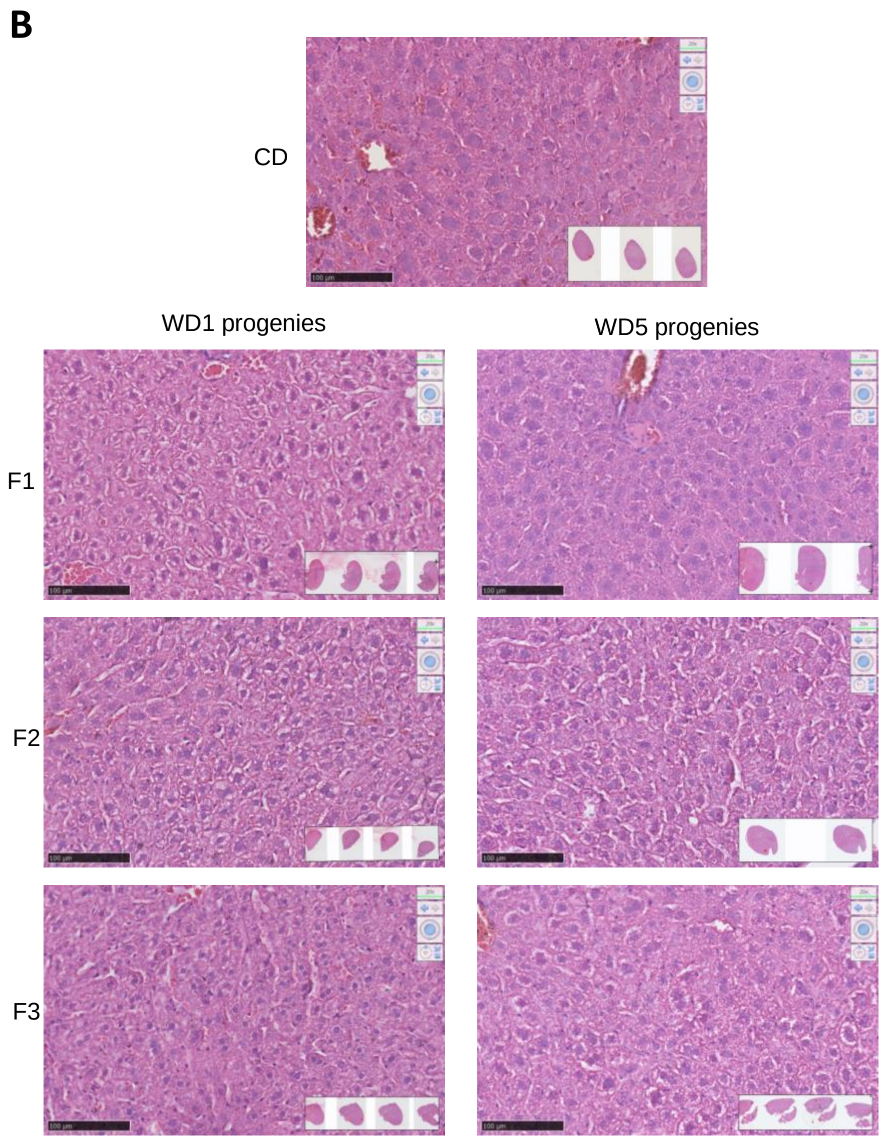
B
CD
WD1 progenies
WD5 progenies
F1
F2
F3

S3 Fig

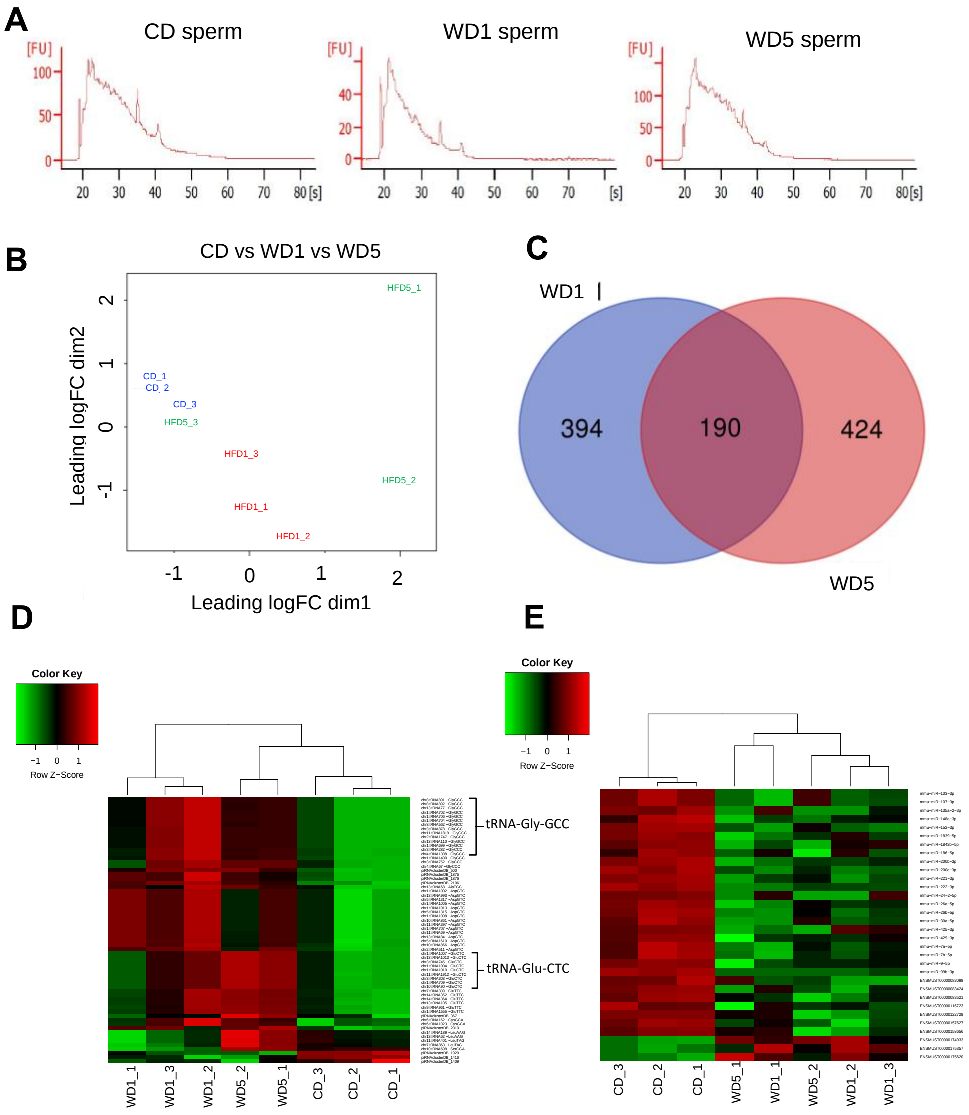
A
CD sperm
WD1 sperm
WD5 sperm
[FU]
[s]
B
CD vs WD1 vs WD5
Leading logFC dim2
Leading logFC dim1
CD_1
CD_2
CD_3
HFD5_1
HFD5_2
HFD5_3
HFD1_1
HFD1_2
HFD1_3
C
WD1
WD5
394
190
424
D
Color Key
Row Z−Score
tRNA-Gly-GCC
tRNA-Glu-CTC
WD1_1
WD1_3
WD1_2
WD5_2
WD5_1
CD_3
CD_2
CD_1
E
Color Key
Row Z−Score
CD_3
CD_2
CD_1
WD5_1
WD1_1
WD5_2
WD1_2
WD1_3

S4 Fig

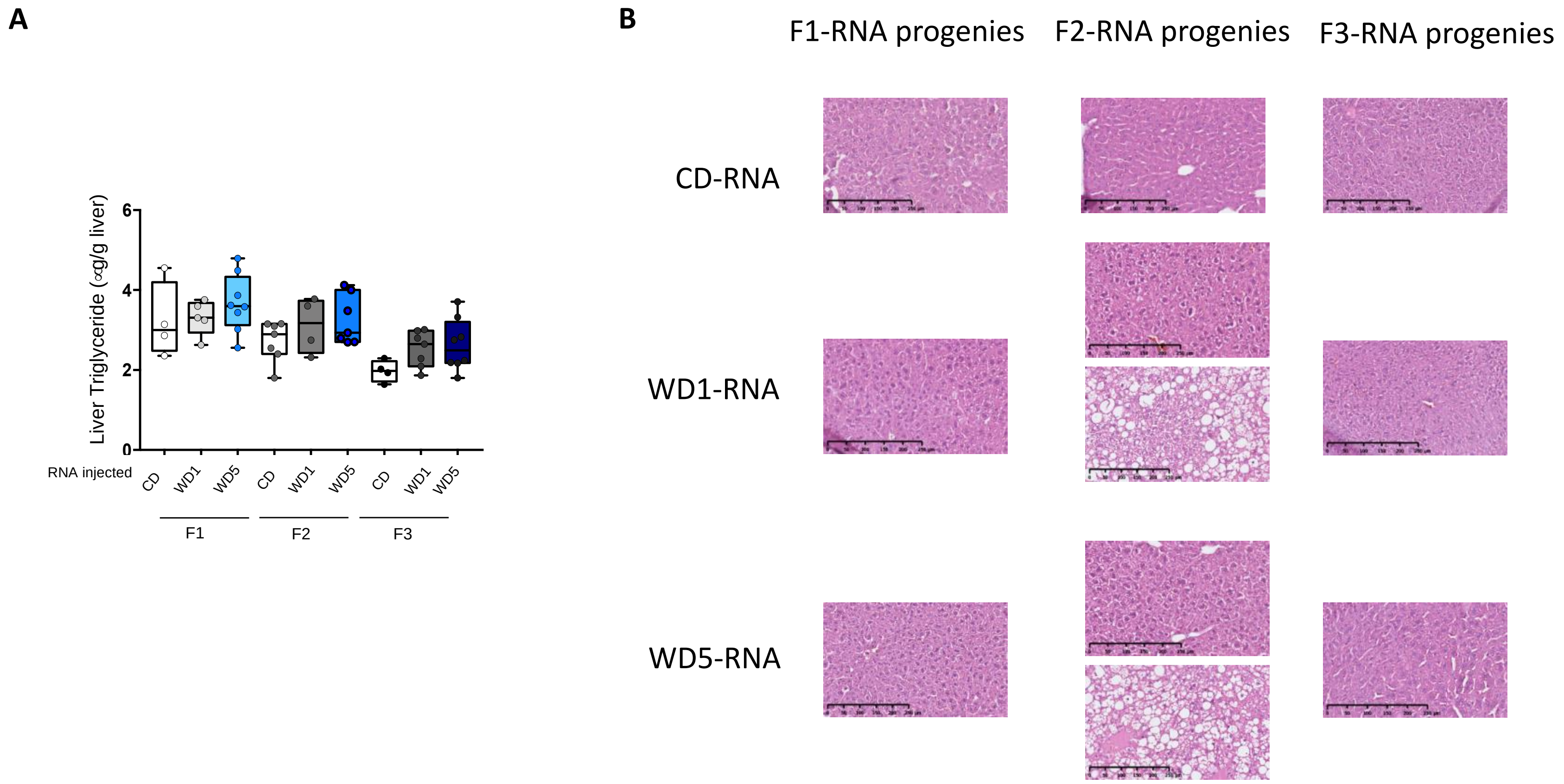
A
Liver Triglyceride (∝g/g liver)
0
2
4
6
RNA injected
CD
WD1
WD5
CD
WD1
WD5
CD
WD1
WD5
F1
F2
F3
B
F1-RNA progenies
F2-RNA progenies
F3-RNA progenies
CD-RNA
WD1-RNA
WD5-RNA

S5 Fig
